# Human MeCP2 binds to promoters and inhibits transcription in an unmethylated *S. cerevisiae* genome

**DOI:** 10.1101/2024.08.12.607623

**Authors:** Joshua A. R. Brown, Maggie Y. M. Ling, Juan Ausió, LeAnn J. Howe

## Abstract

MeCP2 is a DNA-binding transcriptional regulator that is present at near-histone levels in mammalian cortical neurons. Originally identified as a DNA methylation reader, MeCP2 has been proposed to repress transcription by recruiting corepressors to methylated DNA. While some genome-wide occupancy studies support a preference for methylated DNA, others suggest that MeCP2 binding is more influenced by DNA sequence and accessibility than methylation status. Moreover, multiple studies also suggest a role for MeCP2 in gene activation. To clarify MeCP2 function we expressed MeCP2 in *Saccharomyces cerevisiae*, which lacks DNA methylation and known MeCP2 corepressors. We find that MeCP2 is toxic to yeast and globally inhibits transcription, indicating that MeCP2 can have significant functional impacts without DNA methylation or mammalian corepressors. A subset of *MeCP2* mutations that cause the neurodevelopmental disorder Rett syndrome, particularly those that map to the DNA binding domain, alleviate the toxicity of MeCP2 in yeast. Consistent with the importance of DNA binding for toxicity in yeast, we show that MeCP2 binds to the yeast genome, with increased occupancy at GC-rich, nucleosome-depleted sequences. These findings present yeast as a useful tool for analyzing MeCP2 and reveal MeCP2 properties that are not strictly dependent on DNA methylation or mammalian corepressors.

**Summary:** MeCP2, a transcription regulator found in vertebrates, is proposed to repress transcription by recruiting corepressors to methylated DNA. In this work Brown et al. show that MeCP2 expressed in *Saccharomyces cerevisiae*, which lacks DNA methylation and mammalian co-repressors, binds promoters and inhibits transcription. A subset of MeCP2 mutations that cause the neurodevelopmental disorder, Rett syndrome, rescues MeCP2-induced phenotypes in yeast, supporting the relevance of these results to MeCP2 function in mammals.

## Introduction

Rett syndrome (RTT) is a rare neurodevelopmental disorder with limited treatment options, affecting approximately 5-10 of 100 000 females (Petriti et al. 2023). In over 95% of typical RTT cases, a mutation is present in the X-linked methyl-CpG-binding protein 2 (*MeCP2*) gene(Neul et al. 2014; Bin Akhtar et al. 2022), with reoccurring *MeCP2* mutations associated with clinical severity (Neul et al. 2008). Gain-of-function mutation in *MeCP2* can cause MeCP2 duplication syndrome (Ta et al. 2022), reflecting the importance of MeCP2 dosage (Ramocki and Zoghbi 2008), and MeCP2 is increasingly associated with a variety of other disorders including autism and cancer (Ramocki et al. 2009; Kalani et al. 2023; Nejati-Koshki et al. 2023). MeCP2 is approximately 60% unstructured, although specific MeCP2 domains have been defined and characterized, including the highly conserved methyl-binding domain (MBD)(Ghosh et al. 2010). Nearly half of RTT-linked mutations occur within the MBD, including a T158M mutation that is the most common missense mutation in RTT patients (Ghosh et al. 2008; Neul et al. 2008). MeCP2 is found in two isoforms, MeCP2-E1 and MeCP2-E2, which differ only in a short region at the N-terminus. MeCP2-E1 and MeCP2-E2 differ in their physical interactions (Martínez De Paz et al. 2019), but the functional differences between the isoforms are yet to be fully resolved. MeCP2-E2 is only found in mammals (Mnatzakanian et al. 2004) suggesting that the E1 variant was the ancestral vertebrate form of the protein (Martínez De Paz et al. 2019). The E1 variant is the most abundantly expressed in most tissues both in humans (Mnatzakanian et al. 2004) and mice (Kriaucionis and Bird 2004) and mouse *MeCP2* knockouts specific for each variant display an RTT-like phenotype for the E1 knockout only, suggesting that E2 does not functionally compensate for the lack of E1 (Quenard et al. 2006; Fichou et al. 2009). However, as the E2 variant was studied first (Lewis et al. 1992), amino acid numbers follow the common practice of referring to positions in MeCP2-E2.

In addition to its role in RTT, MeCP2 has been extensively studied as a reader of DNA methylation (DNAm) (Good et al. 2021; Bin Akhtar et al. 2022). The MeCP2 MBD shows a preference for methylated DNA in vitro, exhibiting an approximately 3-fold increased binding affinity over unmethylated DNA in a capillary electrophoretic mobility shift assay (Fraga et al. 2003). Additionally, methylated DNA stimulates MeCP2 droplet formation *in vitro*, albeit only to a slightly greater extent than unmethylated DNA (Li et al. 2020). In vivo studies are highly conflicting, with numerous studies supporting both a requirement (Skene et al. 2010; Baubec et al. 2013; Chen et al. 2015; Lagger et al. 2017; Piccolo et al. 2019) and dispensability (Yasui et al. 2007; Rube et al. 2016; Liu et al. 2024) for DNA methylation for genome-wide MeCP2 occupancy. Several explanations for this discrepancy exist. First, recent work suggests that MeCP2’s apparent preference for mCpG sites in vivo is explained by its affinity for GC-rich sequences, and that %GC is a greater predictor of MeCP2 binding than mCpG% (Rube et al. 2016). However, these results are complicated by high levels of DNAm in mammalian genomes, and the tendency of mCpGs to mutate over evolutionary timescales (Hanson and Liebl 2022).

Second, disruption of DNA methylation pathways results in relocation of MeCP2 to open chromatin and recent analysis shows that MeCP2 co-localizes with promoters (Baubec et al. 2013; Liu et al. 2024) suggesting that DNA accessibility may impact MeCP2 binding. Finally, a growing body of evidence also suggests MeCP2 interacts with targets other than DNA itself, specifically RNA (Good et al. 2021) and nucleosomes (Rube et al. 2016; Lee et al. 2020; Ortega-Alarcon et al. 2024). Thus, while the biological importance of MeCP2 in regulating neuronal function is evident, the molecular basis for this is unclear.

Once bound to DNA, common models for MeCP2 function involve MeCP2 binding partners that act as cofactors to affect gene expression (Good et al. 2021). MeCP2 contains a NCoR/SMRT-interaction domain (NID) that promotes MeCP2 binding with the TBLR1 subunit of the mammalian NCoR/SMRT complexes, in a manner dependent on MeCP2-R306 (Lyst et al. 2013; Kruusvee et al. 2017). As such, MeCP2 is proposed to repress gene expression via recruitment of corepressor complexes to DNAm sites (Tillotson and Bird 2020). However, mutations within the NID correspond to some of the milder forms of RTT and over 40 putative binding partners have been reported for MeCP2, many of which represent possible cofactors of MeCP2 other than the NCoR/SMRT complexes (Tillotson and Bird 2020). Indeed, recent work has shown a direct interaction with RNA polymerase II that maps to the C-terminal domain of MeCP2 (Liu et al. 2024), but as the domain of MeCP2 that mediates this interaction has not been tightly defined, it is difficult to determine whether loss of this association is a required feature of RTT. As such, the actual downstream events triggered by MeCP2 binding is a continuing subject of debate.

Many aspects of MeCP2 function are unclear, including the factors regulating DNA binding and the co-factors required for function. To shed light on these we have turned to *Saccharomyces cerevisiae,* a well-characterized model organism that lacks DNAm and a homolog of the NCoR/SMRT complexes, and possesses a nucleosome repeat length similar to mammalian neurons (∼165 bp) (Thomas and Furber 1976; Clark et al. 2020). We found that expression of MeCP2 is severely toxic to yeast, which is corroborated in an accompanying manuscript by Chen *et al*. that surveyed the impact of MeCP2 expression in diverse model systems, including yeast, flies, and mammalian cell lines. Further, MeCP2 expression in yeast inhibits bulk transcription, despite the absence of DNAm and mammalian corepressors. MeCP2 expressed in yeast is specifically enriched at nucleosome-depleted regions (NDRs), while RTT mutations in the MBD rescue yeast growth and abrogate the enrichment of MeCP2 at these sites. These findings reveal a DNAm- and corepressor-independent function for MeCP2 in transcription repression.

## Methods

### Yeast strains and plasmids

Yeast strains used in this study are listed in Supplemental Table S1 and were derived from S288C using standard yeast genetic techniques (Ausubel 1987). Plasmids used in this study are listed in Supplemental Table S2. A yeast codon-optimized MECP2-E1 open reading frame was synthesized by Invitrogen and was amplified with primers containing E2 isoform sequences to produce a MECP2-E2 open reading frame. Wild-type and mutant fragments of the *MeCP2* genes were cloned into the *URA3* centromeric plasmid, pRS416 (Sikorski and Hieter 1989) using the NEBuilder HiFi Assembly Mix and introduced into yeast with a ZF4-ER-VP16 construct integrated at the *LEU2* gene. All *S. cerevisiae* strains were propagated on synthetic complete media lacking uracil and leucine to maintain selection for plasmids and markers.

### Statistics and reproducibility

All experiments were performed with three biological replicates, initially grown from independent yeast colonies, with the exception of the ChIP-seq experiment done with two biological replicates (Figure 3A). All statistical tests are unpaired two-tailed Welch’s *t*-test with n = 3 biological replicates. Exact data underlying the charts in Figures 1D, 1F, and 2C are provided in the Source Data file, along with uncropped blots corresponding to Figures 1A and E. For the ChIP-seq experiment (Figures 3A and S7), spike-in read counts and scaling factors are provided in the Source Data file.

**Figure 1.**
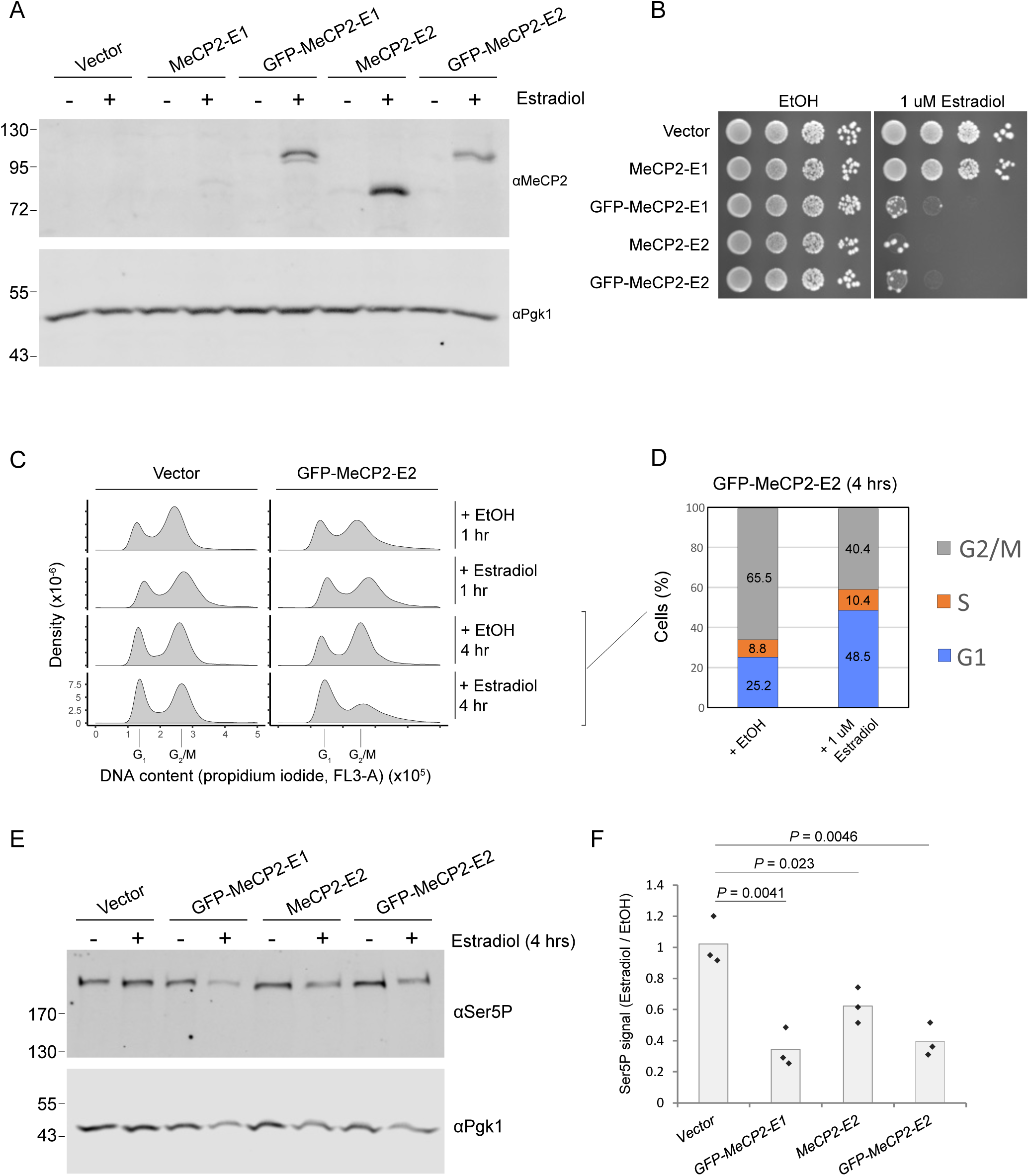
MeCP2 inhibits yeast growth, cell cycle progression, and RNA Pol II transcription. A) Whole-cell extracts are shown from yeast expressing MeCP2 from plasmids, after 1 hour of induction with 1 µM estradiol or treatment with ethanol vehicle. Immunoblotting was performed using anti-MeCP2 antibody, and anti-PGK1 antibody was applied as a loading control. B) 10-fold serial dilutions from overnight cultures were plated on synthetic complete media lacking uracil and leucine to maintain selection for markers. Plates contained 1 µM estradiol to induce MeCP2 expression, or ethanol vehicle, and were grown for three days at 30°C. C) Flow cytometry was performed on fixed yeast cells, stained for DNA content using propidium iodide. Cultures were treated with 1 µM estradiol or ethanol vehicle for 1-4 hours before fixation. A representative sample is shown from three biological replicates. D) Quantification of cell cycle populations from panel B. A mean value scored from the three biological replicates in Source Data is shown. E) Whole-cell extracts were taken from yeast expressing MeCP2 from plasmids, after 4 hours of induction with 1 µM estradiol or treatment with ethanol vehicle. Immunoblotting was performed using anti-Ser5P antibody to measure bulk transcription, and anti-PGK antibody was applied as a loading control. Note that modest decreases in Pgk1 levels were observed in samples expressing MeCP2, consistent with global transcription inhibition. F) Quantification of signal from panel D, bar charts reflect the mean of three biological replicates. Full data are provided in Source Data. All P-values were calculated by two-sided Welch’s t-tests, comparing to wild-type.

### Protein extraction and immunoblotting

For whole cell extracts, overnight cultures were diluted to OD_600_ = 0.2 in synthetic complete media lacking uracil and grown to mid-log phase (OD_600_ = 0.8). Cultures were treated with either 1 µM β-estradiol (Sigma-Aldrich) suspended in ethanol, or the same volume of ethanol as a negative control. Whole cell extracts were prepared as previously described (Kushnirov 2000) and analyzed by SDS-page. Nitrocellulose blots were probed with anti-MeCP2 antibody (Sigma-Aldrich cat # M9317) or anti Ser5P antibody (Millipore cat # 04-1572), and anti-Pgk1 antibody (Invitrogen cat # 459250) was used as a loading control. Blots were visualized using a Biorad ChemiDoc.

### Growth assays

Overnight cultures grown in synthetic complete media lacking uracil were diluted to 0.5 OD_600_, ten-fold serially diluted, and plated on synthetic complete media lacking uracil and leucine containing estradiol. Plates were grown for 3 days at 30°C.

### Flow cytometry

To analyze DNA content, cultures were grown as described for protein extraction. Cells were fixed in 70% ethanol and stored at 4°C. Fixed cells were rehydrated in phosphate-buffered saline, incubated with 100 µg/mL RNase A (ThermoFisher) for 3 hrs, and stained with 5 ug/mL propidium iodide (Invitrogen) for 1 hour. Cells were analyzed using a Cytoflex LX Analyser with gating as described in Figure S1A. Three biological replicates were used to quantify cell cycle populations according to the proportion of cells within each peak in histograms.

To analyze GFP-MeCP2 levels, cultures were grown as described for protein extraction experiments, and induced with 1 μM estradiol for 1 hour. Live cells were analyzed using a Guava easyCyte™ flow cytometer, with gating as described in Figure S5A. Mean signal for three biological replicates is shown in Figure 2C, each full replicate is shown in an empirical cumulative distribution plot in Figure S5B.

**Figure 2.**
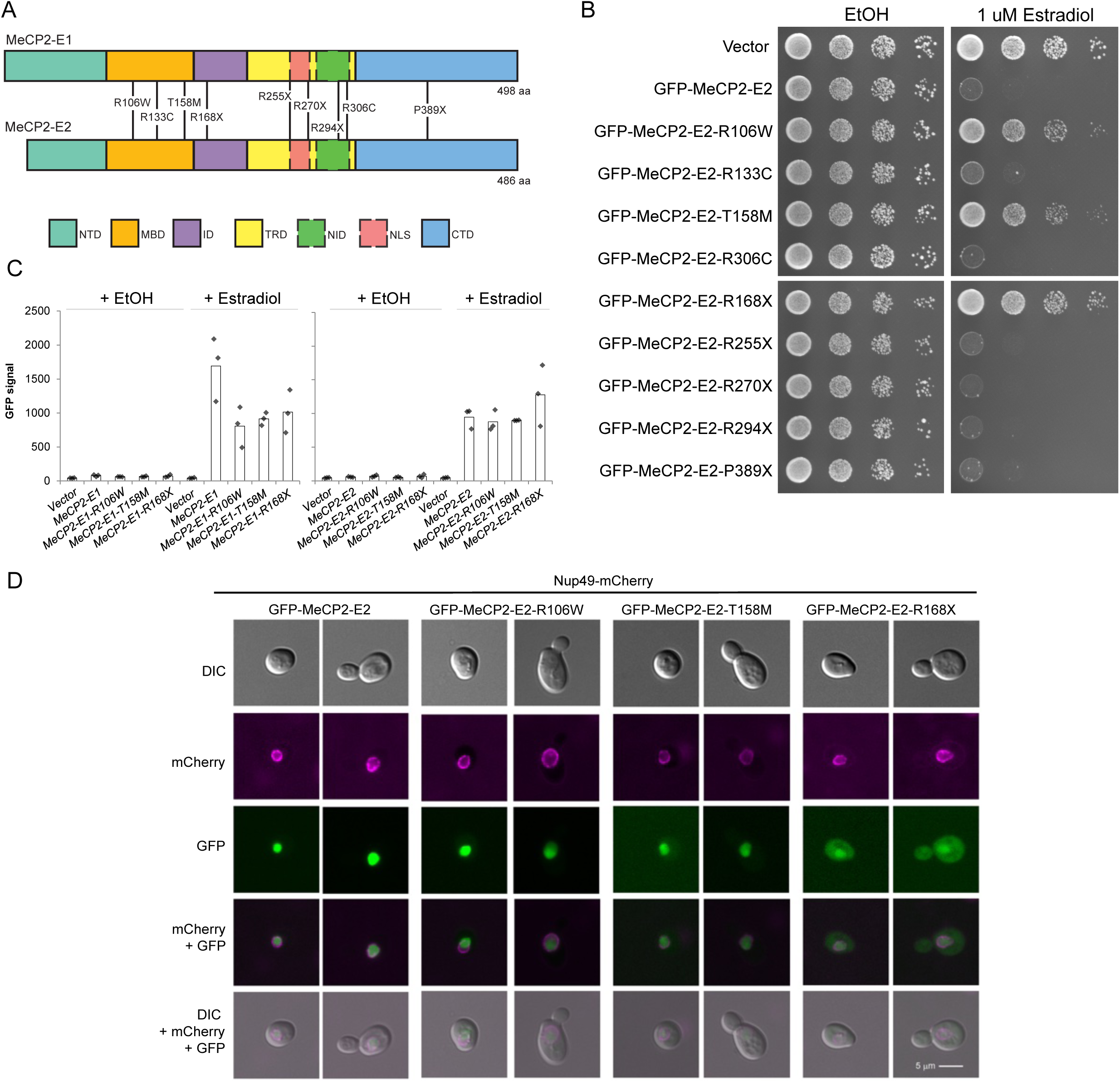
RTT mutations suppress the toxicity of MeCP2 in yeast. A) Diagram of RTT mutations examined in this study, positioned on MeCP2-E1 and MeCP2-E2. Boxes show the positions of annotated MeCP2 features. NTD: N-terminal domain, MBD: methyl-CpG-binding domain, ID: intervening domain, TRD: transcription repression domain, NLS: nuclear localization signal, NID: NCoR/SMRT-interaction domain, CTD: C-terminal domain. Note that the NLS and NID are located within the TRD. B) 10-fold serial dilutions from overnight cultures were plated on synthetic complete media lacking uracil and leucine to maintain selection for markers. Plates contained 1 µM estradiol to induce MeCP2 expression, or ethanol vehicle, and were grown for three days at 30°C. C) Quantified mean signal of three biological replicates of flow cytometry against the GFP tag in yeast cells expressing GFP-MeCP2-E1 or GFP-MeCP2-E2 for 1 hour. Full data are provided in the Source Data file. D) Representative cells are shown from fluorescent microscopy of yeast expressing GFP-MeCP2-E2 (green) variants for one hour, while constitutively expressing Nup49-mCherry (magenta) as a marker of the nuclear membrane. A differential interference contrast (DIC) image is also shown.

### Microscopy

Cultures were grown to mid-log and treated with 1 μM estradiol for 1 hour to induce MeCP2 expression. Live cells were spun, resuspended in 1X PBS, and mounted on concanavalin A-coated slides. Microscopy was performed using a Leica THUNDER 3D Cell Imager with Leica K8 CMOS camera using a 100x (oil immersion) objective (HC PL APO). Images were captured at a resolution of 2048×2048 pixels, with a pixel size of 15.2881 pixels per micron.

Images were analyzed using ImageJ. Eleven images were obtained at 0.3 micron intervals along the z-axis, and localization was reviewed across all focal planes. A representative budded and unbudded cell is shown for each strain, as all cells with clear fluorescent signals showed the same localization pattern.

### ChIP sequencing

Budding yeast were diluted from overnight cultures into synthetic complete media lacking uracil and grown to mid-log phase at 30°C before treatment with 1 µM estradiol for one hour. At a final OD_600_ of 0.8, cultures were crosslinked with 1% formaldehyde for 20 min, quenched with 300 mM glycine for 5 min, and frozen at −70°C. For each sample, 450 million budding yeast cells were combined with 68 million wild-type *Schizosaccharomyces pombe* cells, as a spike-in, which had been crosslinked and frozen by the same approach. Cells were resuspended in 0.6 mL of lysis buffer (50 mM Hepes-KOH [pH 7.5], 140 mM NaCl, 1 mM EDTA, 1% Triton X-100, 0.1% sodium deoxycholate) and lysed by bead beating before centrifugation, then resuspended in 1 mL. Samples were sonicated with a Covaris S220 focused-ultrasonicator (peak power 150 W for 7 min with 30 s cycling off/on, duty factor 14, 200 cycles/burst). One hundred microlitres of lysate was reserved as input and 750 uL was used for immunoprecipitation. Samples were precleared with protein G Dynabeads™ (ThermoFisher cat# 10003D) for 1 hour, incubated overnight with anti-GFP antibody (Abcam cat# ab290), and incubated with protein G beads for 1 hour, all at 4°C. Beads were washed twice with lysis buffer (50 mM HEPES pH 7.5, 140 mM NaCl, 0.5 mM EDTA, 1% triton X-100, 0.1% sodium deoxycholate), twice with lysis buffer with 500 mM NaCl, and twice with LiCl wash buffer (10 mM Tris-HCl pH 8.0, 250 mM LiCl, 0.6% NP-40, 0.5% sodium deoxycholate, 1 mM EDTA), before 1 wash with TE (pH 8). Samples were eluted in elution buffer (TE pH 8, 1% sodium dodecyl sulfate, 150 mM NaCl, 5 mM dithiothreitol) three times by incubation at 65°C for ten minutes, with shaking.

Sequencing libraries were prepared as described previously (Maltby et al. 2012). Briefly, DNA was end-repaired, A-tailed, and ligated to adapters before 17 cycles of PCR amplification with indexed primers. All DNA purification steps used Solid Phase Reversible Immobilization magnetic beads (Omega, M1378). Samples were pooled, gel-purified (200-600 bp fragments) and sequenced on an Illumina NextSeq.

### ChIP-seq analysis

Sequence reads were trimmed for adapter sequences using cutadapt (http://cutadapt.readthedocs.io/en/stable/) and mapped to the Saccer3 genome build using BWA-MEM. Data was analyzed using DeepTools (Ramírez et al. 2016) using the Galaxy web platform (Afgan et al. 2016). bamCoverage was used to analyze IP and input samples separately, while bamCompare was used to compare IP and input signal. Parameters used were: bin size = 5 bp, mapping quality threshold >= 25, paired-end read extension, duplicate reads removed. Only reads mapped in proper pairs by BWA-MEM were used in subsequent analysis.

IP and input samples were normalized based on the relative number of reads mapping to the genome of the *S. pombe* spike-in. “IP/input” analyses are presented as the log_2_ of IP signal divided by input signal, in which IP reads were multiplied by a normalization factor (1000 / *S. pombe* IP reads), as in the Source Data file. This value of 1000 was used as an arbitrary scaling factor while dividing by *S. pombe* spike-in reads, similar to the use of scaling factors in previous work (Jeronimo et al. 2019). Input samples were multiplied by a normalization factor of (1 / *S. pombe* input reads). Heatmaps of separate IP and input samples (Figure S7) show the same analysis, with IP and input samples separately divided by respective spike-in reads. Input scaling was performed using *S. pombe* spike-in in this study to account for the possibility of varying input amounts, as yeast cells expressed toxic MeCP2 in these ChIP experiments. MeCP2 expression could affect the cell cycle status of yeast, thereby altering the amount of input DNA extracted from each cell used for ChIP. Note that our results appeared similar with or without adjusting input by spike-in in this manner.

Heatmaps represent distance from the position of the +1 nucleosome (Chereji et al. 2018), and genes were sorted according to the length of their NDR (Chereji et al. 2018). Previously-published MNase-seq data was shown alongside select heatmaps as a visual aid for NDRs (Martin et al. 2021). Subsequent analyses using quantiles were performed using Java Genomics Toolkit (https://github.com/timpalpant/ java-genomics-toolkit).

### Boxplots

All boxplots used to display ChIP-seq analysis extend from the first to third quartiles. Whiskers extend to the most extreme data points, or to 1.5 times the interquartile range from the box, whichever is lesser. Notches represent +/-1.58 interquartile range/sqrt(n), and non-overlapping notches can approximate a 95% confidence interval for comparing medians.

## Results

### MeCP2 inhibits yeast growth, cell cycle progression, and RNA Pol II transcription

We sought to directly interrogate whether MeCP2 expression in yeast could impact gene expression despite the absence of DNAm or the NCoR/SMRT complexes. To do so, we made use of budding yeast strains containing centromeric plasmids with MeCP2 expression controlled via estradiol (see Methods). We first examined the stability of MeCP2 isoforms, with and without an N-terminal sfGFP tag. MeCP2 was detected after 1 hour of induction with estradiol (Figure 1A), indicating that budding yeast were suitable for controlled MeCP2 expression. An N-terminal GFP tag was necessary for clear detection of the MeCP2-E1 variant, but the MeCP2-E2 variant did not require the tag for stable expression. Yeast expressing GFP-MeCP2-E1, MeCP2-E2, and GFP-MeCP2-E2 exhibited a significant growth defect (Figure 1B), but the untagged MeCP2-E1 construct did not cause a similar effect, consistent with the low level of MeCP2-E1 in cells (Figure 1A). These results confirm that MeCP2 can be expressed in yeast, and significantly impacts growth, while the low level of untagged MeCP2-E1 suggests that the E1 variant is unstable without an N-terminal GFP tag.

We next sought to determine how MeCP2 inhibited yeast growth. We first examined progression through the yeast cell cycle by flow cytometry of cultures expressing GFP-MeCP2-E2 for 1-4 hours. DNA staining revealed an increase in the G_1_ phase population after 4 hours of MeCP2 induction via estradiol, along with a decrease in G_2_/M phase cells (Figures 1C, D, and S1B). We hypothesized that MeCP2 delays cell-cycle processes by inhibiting transcription, despite the absence of DNAm or the NCoR/SMRT complexes and examined bulk RNA Pol II transcription activity in cells expressing MeCP2 for four hours (Figures 1E and F). GFP-MeCP2-E1, MeCP2-E2, and GFP-MeCP2-E2 each reduced the level of RNA Pol II serine 5 phosphorylation (Ser5P), a mark of global transcription initiation. This inhibition of bulk transcription initiation may contribute to the cell cycle delay we observed in yeast expressing MeCP2. Collectively, these data indicated that human MeCP2 has significant impacts on yeast growth and transcription, despite the absence of DNAm or the mammalian NCoR/SMRT complexes.

### RTT mutations suppress the toxicity of MeCP2 in yeast

The toxicity of MeCP2 in yeast suggested that yeast may be a useful platform for determining whether common RTT mutations have significant impacts on DNAm- and corepressor-independent MeCP2 activity. We therefore used yeast to express the most common MeCP2 mutations underlying RTT, collectively accounting for approximately half of RTT cases (Figure 2A) (Krishnaraj et al. 2017). We found that the R106W, T158M, and R168X mutations largely suppressed the toxicity of MeCP2 in yeast, whether in the context of GFP-MeCP2-E2 (Figure 2B), MeCP-E2 (Figure S2), or GFP-MeCP2-E1 (Figure S3). In contrast, mutations that disrupted the interaction of MeCP2 with the NCoR/SMRT complexes (R306C) or with RNA Pol II (R270X) did not abolish toxicity (Lyst et al. 2013; Liu et al. 2024). We considered that high MeCP2 induction with 1 μM estradiol could mask intermediate phenotypes, and therefore also examined RTT mutations while expressing GFP-MeCP2-E2 with 50 nM estradiol, which revealed a partial suppression of toxicity by the R133C and R255X mutations (Figure S4). To rule out the possibility that the most strongly-suppressing mutations reduced the stability or nuclear localization of MeCP2, we first quantified protein levels by GFP flow cytometry, allowing us to determine the consistency of protein levels across thousands of cells in our tagged constructs. We found no clear evidence that the stability of GFP-MeCP2-E1 or GFP-MeCP2-E2 was greatly altered by the R106W, T158M, or R168X mutations, in the context of MeCP2 induction by 1 μM estradiol (Figures 2C and S5). We then examined protein localization by fluorescent microscopy, comparing the GFP tag signal to Nuclear Pore 49 (Nup49) tagged with mCherry as a marker of the nuclear membrane. Using this system, we confirmed that the R106W and T158M mutations did not disrupt nuclear localization, but found that the R168X mutation caused a cytoplasmic dispersal of GFP-MeCP2-E2 (Figure 2D) and GFP-MeCP2-E1 (Figure S6). Because the R106W and T158M mutations map to the DNA-binding domain of MeCP2, this suggests that growth inhibition by MeCP2 is dependent on its DNA-binding ability.

### MeCP2 is enriched at NDRs in the yeast genome, in a T158-dependent manner

Our findings up to this point suggest that MeCP2 inhibits transcription in yeast and this is dependent on its DNA binding ability. To confirm this, we used ChIP-seq to determine if MeCP2 bound to the unmethylated yeast genome. We examined genome-wide occupancy of GFP-MeCP2-E2 using an anti-GFP antibody, to enable comparison to a stable untagged MeCP2-E2 control. We also included the T158M mutation, which is the most common RTT missense mutation and affects DNA affinity (Ghosh et al. 2008; Neul et al. 2008; Yang et al. 2016; Good et al. 2021). Consistent with previously published works using mammalian cells (Thambirajah et al. 2012; Lee et al. 2020), ChIP of GFP-MeCP2-E2 revealed a striking enrichment at NDRs, as exemplified in heatmaps ordered by the size of the well-characterized NDRs (Martin et al. 2021) overlapping the promoters of protein coding genes (Figure 3A, two biological replicates shown) (Chereji et al. 2018). MeCP2 enrichment within NDRs was also apparent when viewing heatmaps of IP and input samples separately (Figure S7). This NDR enrichment was abolished by the T158M mutation, suggesting that T158 is required for MeCP2’s affinity for unmethylated DNA, and indicating that NDR enrichment was not simply an artifact of our ChIP-seq method. To clarify the relationship between nucleosome occupancy and MeCP2 enrichment, we sorted the yeast genome into 100 bp intervals and split these into deciles based on nucleosome occupancy data (see Methods). In this analysis, GFP-MeCP2-E2 was enriched in the bottom decile of nucleosome occupancy, compared to the second decile, a trend not observed for the T158M mutant (Figure 3B). This suggests that MeCP2 was specifically enriched at NDRs in an MBD-dependent manner. Our data also indicated that both GFP-MeCP2-E2 and GFP-MeCP2-E2-T158M were present across genes outside of these NDRs, albeit at a lower level in the mutant. This broad distribution is reminiscent with what is observed in mammalian cells, with MeCP2 bound to large fractions of the genome (Lagger et al. 2017).

**Figure 3.**
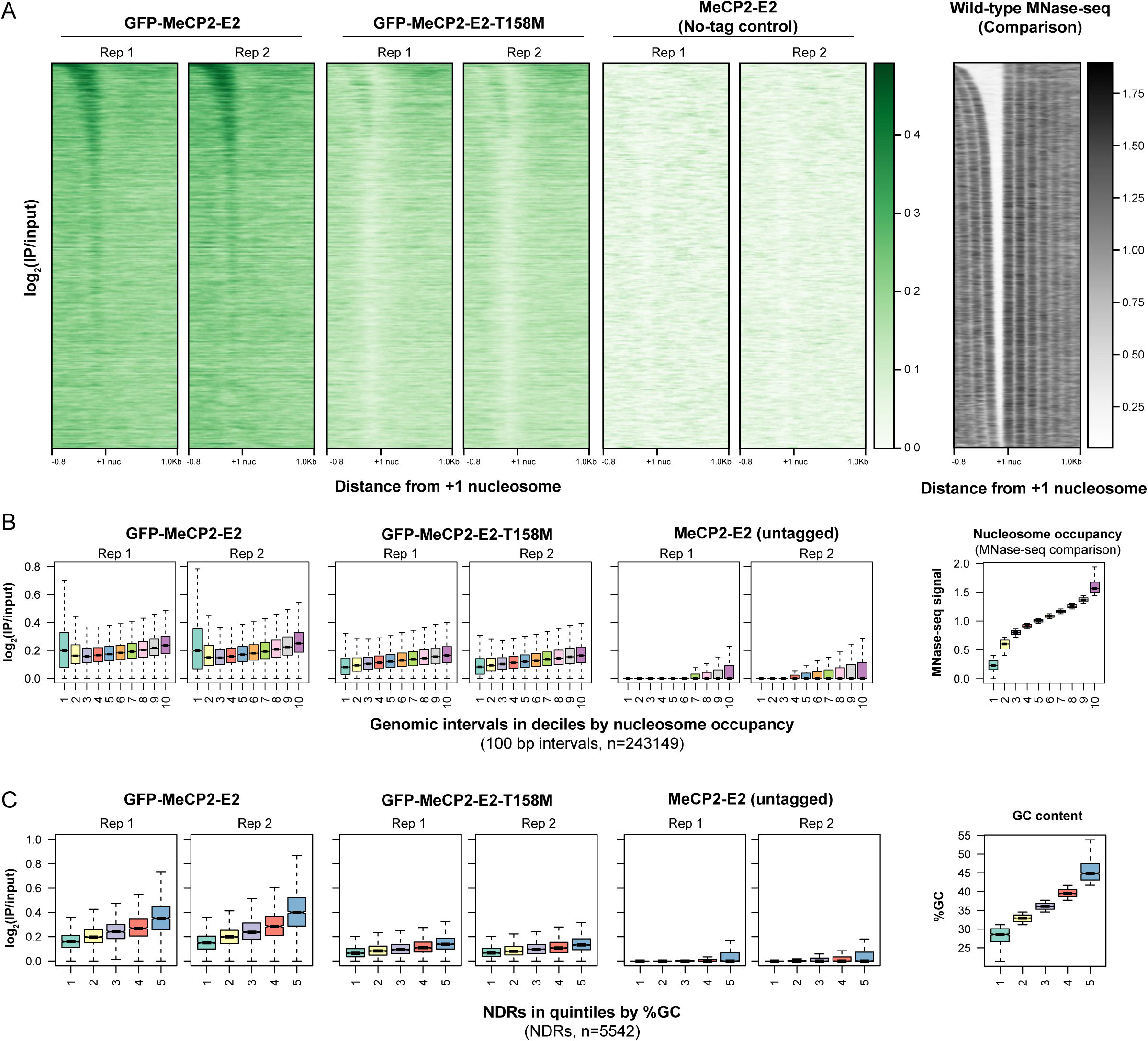
MeCP2 is enriched at NDRs in the yeast genome, in a T158-dependent manner. A) Heatmaps of ChIP-seq performed using anti-GFP antibody, on yeast cells induced with estradiol to express GFP-tagged MeCP2 for one hour. The x-axis represents distance from the +1 nucleosome ^46^. Heatmap signal represents log2(IP/input) signal, after normalization according to spike-in reads (see Methods). A wild-type MNase-seq experiment showing nucleosome positions is displayed alongside the ChIP-seq experiment for comparison purposes, and all genes are ordered according to the size of the NDR ^46^. B) The yeast genome was divided into 243149 intervals of 100 bp in length with a 50 bp sliding window, and sorted into deciles by nucleosome occupancy in previously-published MNase-seq data. ChIP-seq enrichment of MeCP2, as in panel A, is shown for each interval in a decile. Boxplots are centered on median signal as described in Methods. C) MeCP2 ChIP-seq signal within NDRs correlated with %GC. 5542 NDRs ^46^ were divided into quintiles based on %GC. The GFP-MeCP2-E2 ChIP-seq data in panel A was analyzed for enrichment in each NDR in each bin. Boxplots are centered on median signal as described in Methods.

Next, we took advantage of our ChIP-seq data to interrogate the claim that MeCP2 binding can be driven by GC content (Rube et al. 2016), again taking advantage of the absence of DNAm which might confound such analysis. We sorted each NDR in the yeast genome (Chereji et al. 2018) into quintiles based on GC content. According to this analysis, GFP-MeCP2 enrichment was greater at NDRs with higher GC content (Figure 3C). We then tested the possibility that this enrichment was caused by a correlation between GC content and NDR size. To do so, we split each NDR into quintiles based on NDR width (Chereji et al. 2018) before splitting each quintile into five additional quintiles based on %GC. This analysis revealed that GFP-MeCP2-E2 signal was increased at NDRs with a higher %GC, even within deciles set by NDR length (Figure S8).

Together, these findings indicate broad binding of MeCP2 to the unmethylated yeast genome, along with a specific enrichment at NDR sequences, particularly ones with higher GC content. This reveals MeCP2 preferences for DNA binding in vivo in the absence of DNAm pathways.

## Discussion

In this study we show that MeCP2 expression inhibits growth and transcription in budding yeast. Genome-wide profiling of wild-type and mutant MeCP2 occupancy suggests that these phenotypes are likely a consequence of MeCP2 binding to promoters, revealing DNAm-independent targeting of MeCP2 in vivo and repression in the absence of the NCoR/SMRT complex. Common RTT mutations in the MBD rescue toxicity and disrupt promoter binding, revealing that these clinically relevant mutations affect MeCP2 properties that are not strictly dependent on genome methylation or recruitment of corepressor complexes.

Based on our results, we propose that DNA binding by MeCP2 in vivo is regulated by chromatin accessibility and not strictly dependent on DNAm. Given MeCP2’s role as a DNAm reader, this result is perhaps unexpected, but our findings do not contradict the large body of prior research regarding MeCP2 function. Rather, our results emphasize that MeCP2’s lower-affinity interactions with unmethylated DNA can have significant biological impacts. This interpretation is consistent with numerous published works. First, in embryonic stem cells, in addition to a correlation with mCpG, MeCP2 was associated with a cluster of active and accessible chromatin marks, and depletion of DNAm increased association with these sites (Baubec et al. 2013). Second, more recent work demonstrates co-localization of MeCP2 and RNA Pol II downstream of TSSs (Liu et al. 2024). MeCP2 binding to unmethylated DNA may also be affected by factors such as sequence, as shown by the data in this study and others (Rube et al. 2016), or MeCP2 abundance. The levels of MeCP2 can vary 7-fold between neural and non-neural brain cells (Skene et al. 2010), and MeCP2 levels in neurons are on the same order of magnitude as mCpGs and nucleosomes (Skene et al. 2010). At these levels, MeCP2 may be able to saturate its highest-affinity mCpG binding sites, leaving sufficient MeCP2 to bind less favourable sites. This may also be the case in conditions such as cancer, MeCP2 duplication syndrome (Kalani et al. 2023), or mouse models of detrimental MeCP2 overexpression (Collins et al. 2004; Luikenhuis et al. 2004). An example of MeCP2 enrichment being changed by cellular context, rather than altered DNAm, would be studies of a peripheral nerve injury model in mice which found higher MeCP2 levels in dorsal root ganglia and differences in MeCP2 binding at a microRNA locus, without a change in DNAm at this locus (Manners et al. 2015; Manners et al. 2016). Finally, the lower levels of MeCP2-E1 in yeast (Figure 1A) suggests that the N-terminus of the E1 variant can confer protein instability in certain conditions, which could exacerbate the effect of mutations that affect MeCP2 protein levels.

RTT mutations have diverse impacts on the toxicity of MeCP2 in yeast, and our findings suggest that growth inhibition is dependent on the DNA-binding properties of the MBD. For example, the R106W and T158M mutations effectively suppressed toxicity in yeast, consistent with their impact on DNA binding in vitro, while the R133C mutation only partially suppressed MeCP2 toxicity (Figure S4), reflecting the limited impact of this mutation on MeCP2 binding to unmethylated DNA in vitro (Nikitina et al. 2007). In addition to disrupting DNA binding, the MeCP2-T158M mutation destabilizes MeCP2 in knock in mice, and the resulting RTT-like phenotypes can be partially suppressed by MECP2 overexpression, suggesting an importance of protein levels in this context (Lamonica et al. 2017). Interestingly, however, our findings highlight that the T158M mutation can significantly impact *MeCP2* phenotypes, despite a similar protein level to wild type. The RTT mutations that did not fully restore yeast growth likely affect MeCP2 properties that are inconsequential in yeast, such as binding to cofactors. For example, the MeCP2-R306C mutation is important for interaction with the NCoR/SMRT corepressor complexes (Lyst et al. 2013; Kruusvee et al. 2017), but is largely dispensable for MeCP2 protein stability and DNA binding (Boxer et al. 2020). The lack of an effect of the MeCP2-R306C mutation on yeast growth is consistent with the absence of the NCoR/SMRT complexes in yeast. Notably, an accompanying study by Chen *et al*. recapitulates the lack of an MeCP2-R306C phenotype in yeast, but identifies suppression by R306C in *Drosophila melanogaster*, consistent with an effect on MeCP2’s co-repressors. Similarly, the lack of growth rescue by C-terminal truncations beginning at MeCP2-R270 suggests that cofactors interacting with the C-terminal region of MeCP2 are dispensable for growth inhibition in yeast. This includes RNA Pol II, which has recently been shown to interact with MeCP2 in a manner that is diminished by R270X truncation (Liu et al. 2024).

MeCP2’s role in gene repression is often associated with corepressors such as the NCoR/SMRT complexes (Good et al. 2021), yet the ability of MeCP2 to inhibit transcription in yeast is independent of these factors. It is possible that MeCP2’s binding to exposed promoters interferes with binding of the core transcription machinery (Kaludov and Wolffe 2000). Indeed, a MeCP2 fragment containing the MBD and lacking the NCoR interaction domain weakly represses transcription in vitro (Nan et al. 1997). In addition to binding NDRs, we observe lower levels of MeCP2 bound over gene bodies. Similar genome-wide occupancy has been observed in neurons (Tillotson and Bird 2020), and MeCP2 can bind and compact chromatin in vitro (Nikitina et al. 2007). However, it is unlikely that this property explains the transcription repressive activities of MeCP2 in yeast as chromatin interaction requires the C-terminal domain of MeCP2 (Georgel et al. 2003), which is dispensable for growth inhibition in yeast.

Our findings reveal that MeCP2 can bind to an unmethylated genome and, despite the absence of known corepressors, can negatively impact transcription. Notably, MeCP2 phenotypes in yeast can be substantially affected by certain RTT mutations, particularly those within the MBD. These collective results suggest that DNAm- and corepressor-independent MeCP2 functions may be worth greater consideration in future research of MeCP2 function, and of RTT mutations that affect the MBD.

## Data availability

Strains and plasmids are available upon request. Data generated for this manuscript were deposited in the NCBI Gene Expression Omnibus under the accession code “GSE269155”. Published datasets analyzed for this paper include “GSM4850570” and “GSM4850572.” (*S. cerevisiae* MNase-seq) and https://static-content.springer.com/esm/art%3A10.1186%2Fs13059-018-1398-0/MediaObjects/13059_2018_1398_MOESM2_ESM.xlsx (the coordinates of the dominant positions of +1 and –1 nucleosomes in *S. cerevisiae*).

## Acknowledgements

This work was supported by the LSI Imaging Core Facility (RRID:SCR_023783) and the UBC Flow core facility at the University of British Columbia, supported by Life Sciences Institute, the UBC GREx Biological Resilience Initiative.

---

**Supplementary Figure 1.**
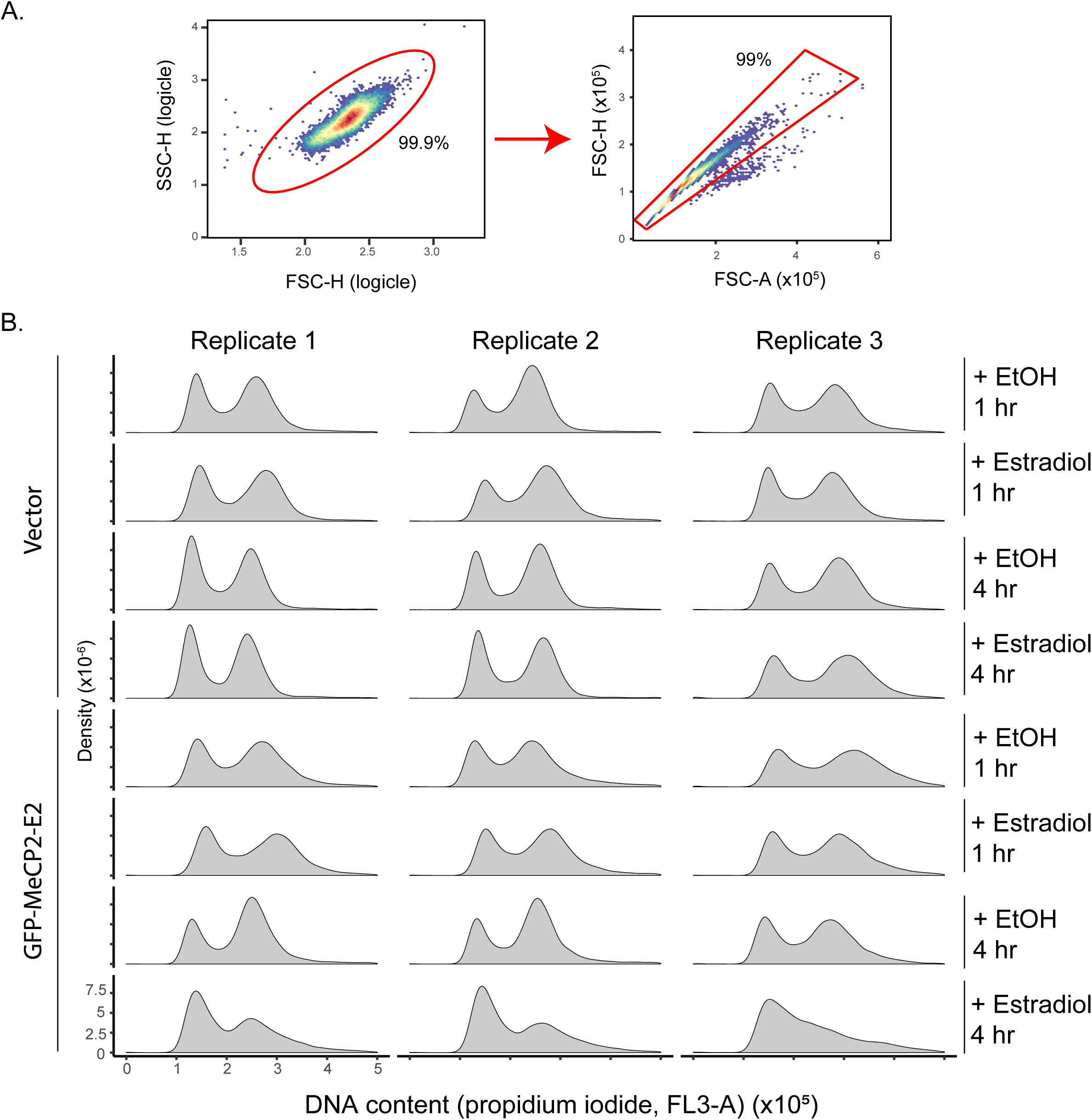
Raw data for flow cytometry on yeast expressing GFP-MeCP2-E2. A) Gating strategy used for cell cycle analysis by flow cytometry after propidium iodide staining. Events were gated by FSC-H vs SSC-H to remove debris, then to remove possible doublet events by FSC-A vs FSC-H. B) Density plots of cell cycle analysis by flow cytometry for each of three biological replicates. Flow cytometry was performed as in Fig. 1c, d. The percentage of cells counted within G1, S, and G2/M peaks for each replicate is provided in Source Data, over twenty thousand cells were counted for each replicate.

**Supplementary Figure 2.**
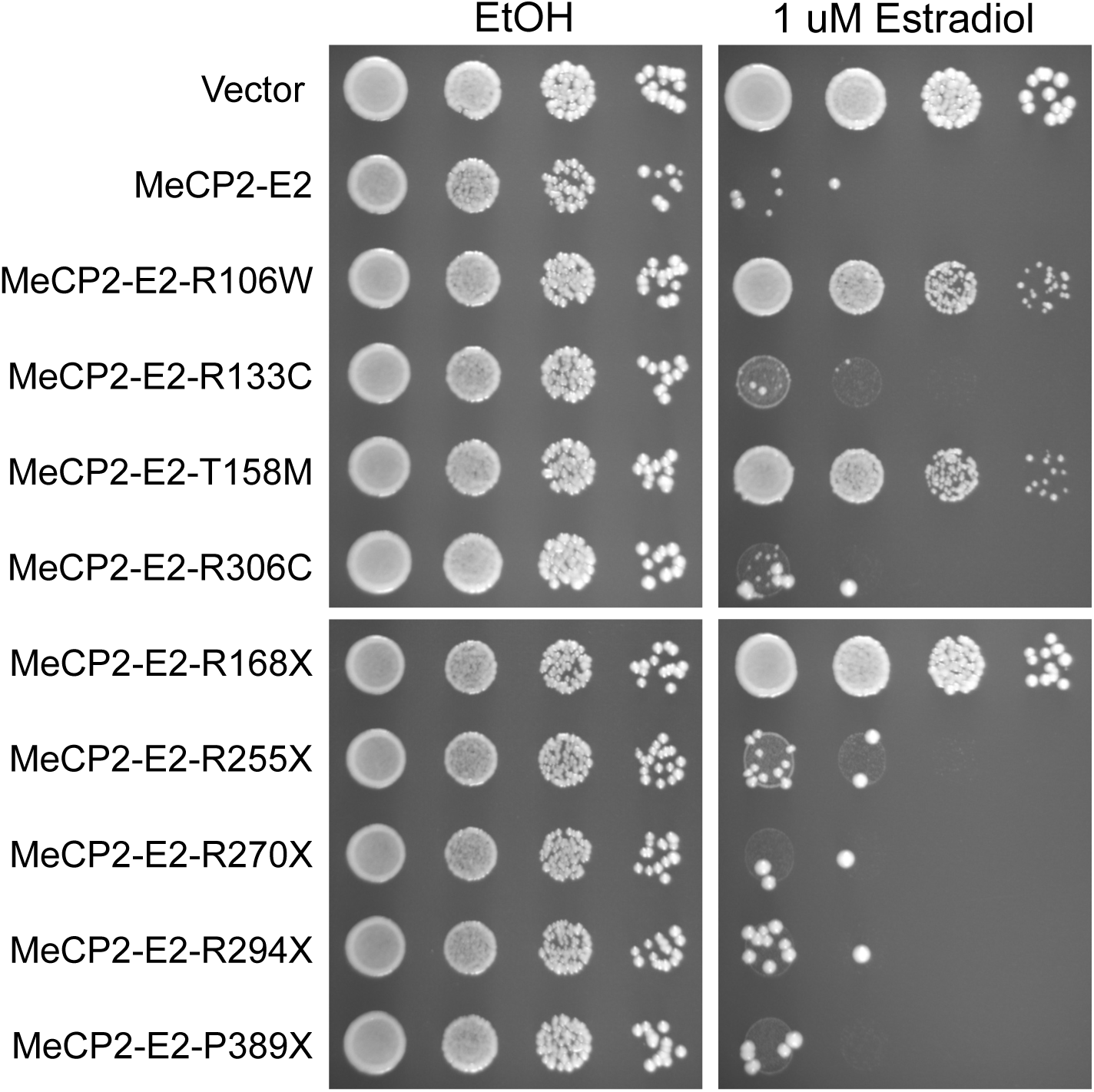
Specific RTT mutations suppressed the toxicity of MeCP2-E2 in yeast. 10-fold serial dilutions from overnight cultures were plated on synthetic complete media lacking uracil and leucine to maintain selection for markers. Plates contained 1 µM estradiol or ethanol vehicle, and were grown for three days at 30°C.

**Supplementary Figure 3.**
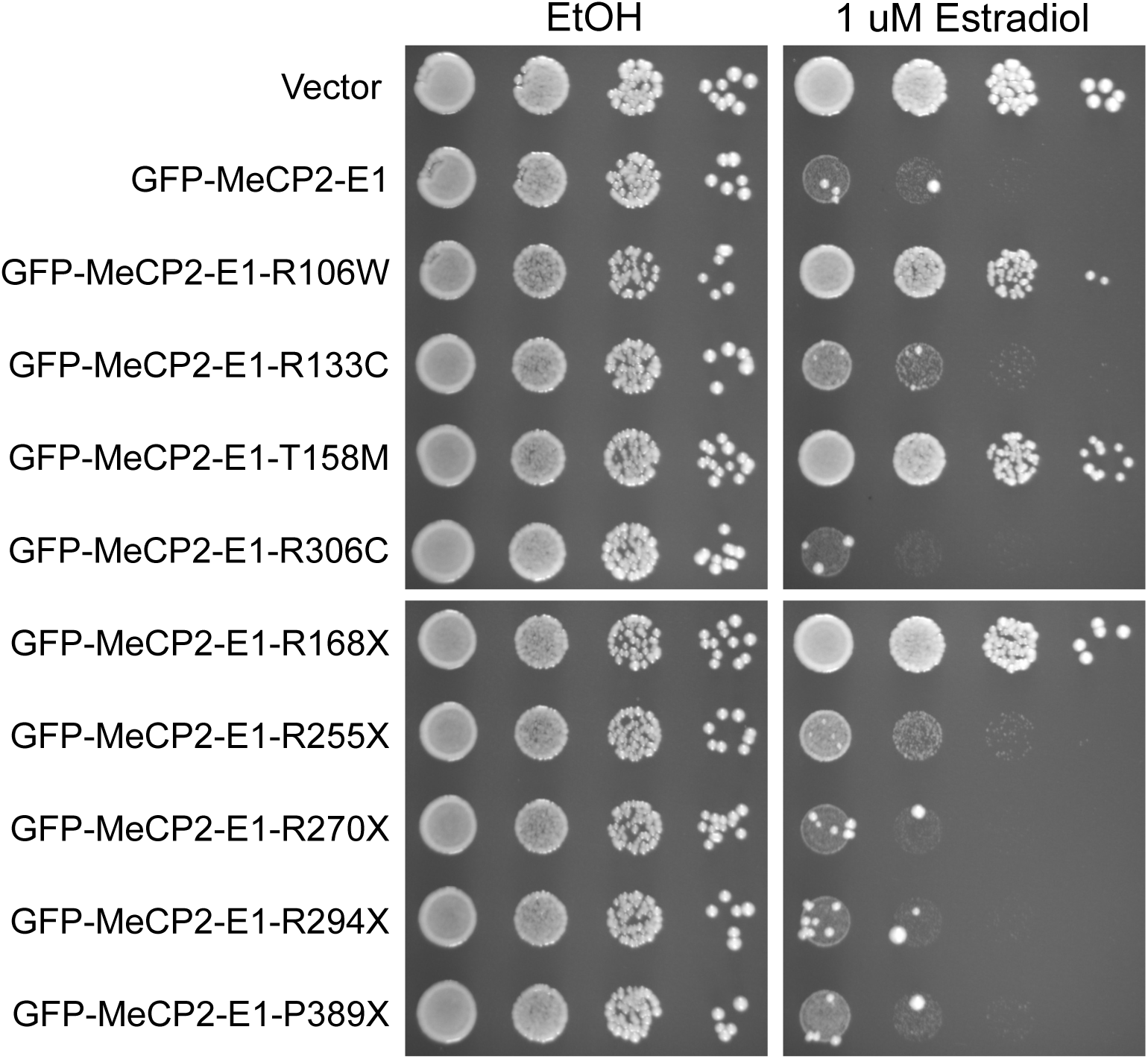
Specific RTT mutations suppressed the toxicity of GFP-MeCP2-E1 in yeast. Plates were grown as in Supplementary Figure 2.

**Supplementary Figure 4.**
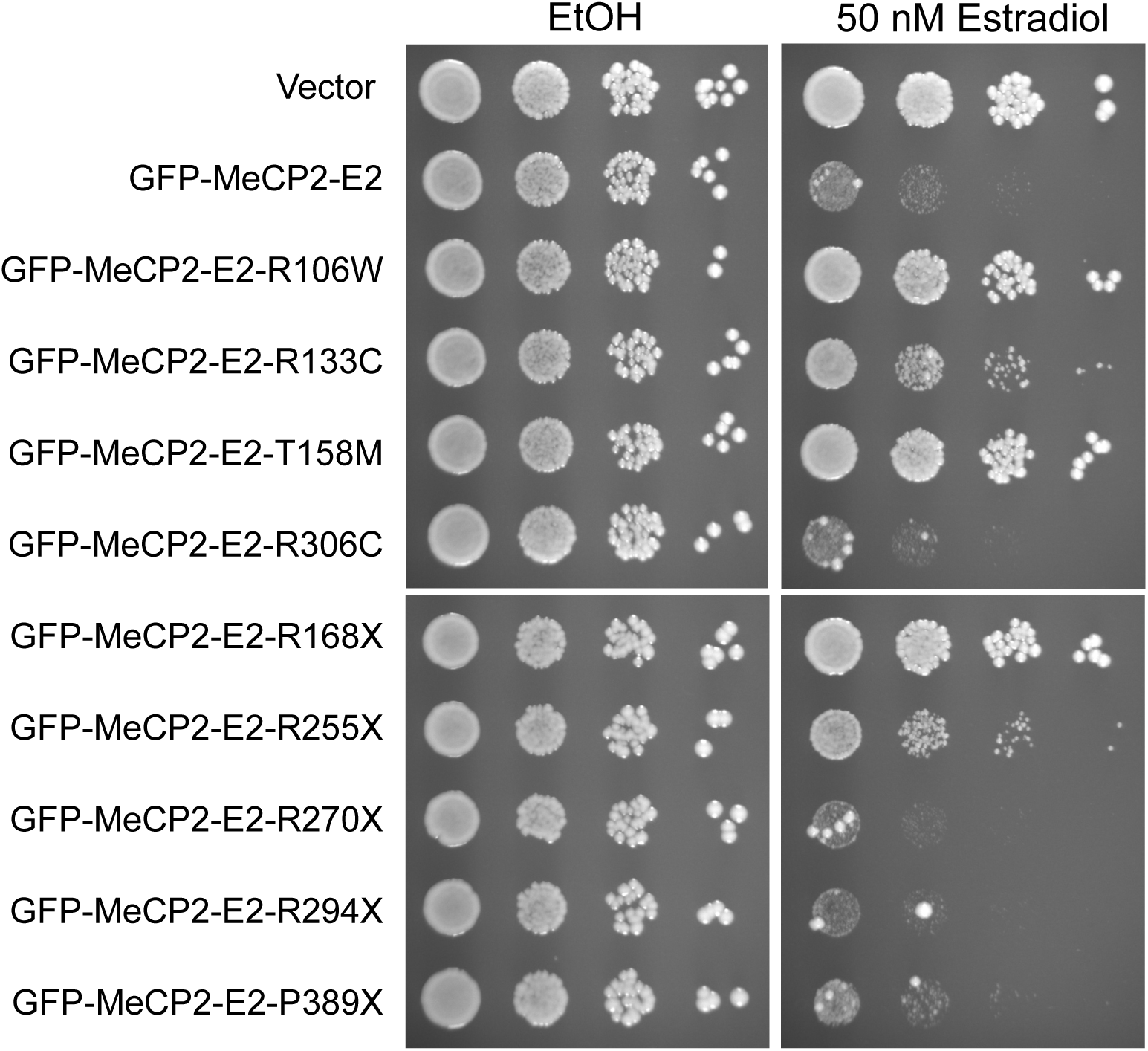
Suppression of GFP-MeCP2-E2 phenotypes by RTT mutations in the context of induction by 50 nM estradiol. Plates were grown as in Supplementary Figure 2, except that 50 nM estradiol was used to induce MeCP2.

**Supplementary Figure 5.**
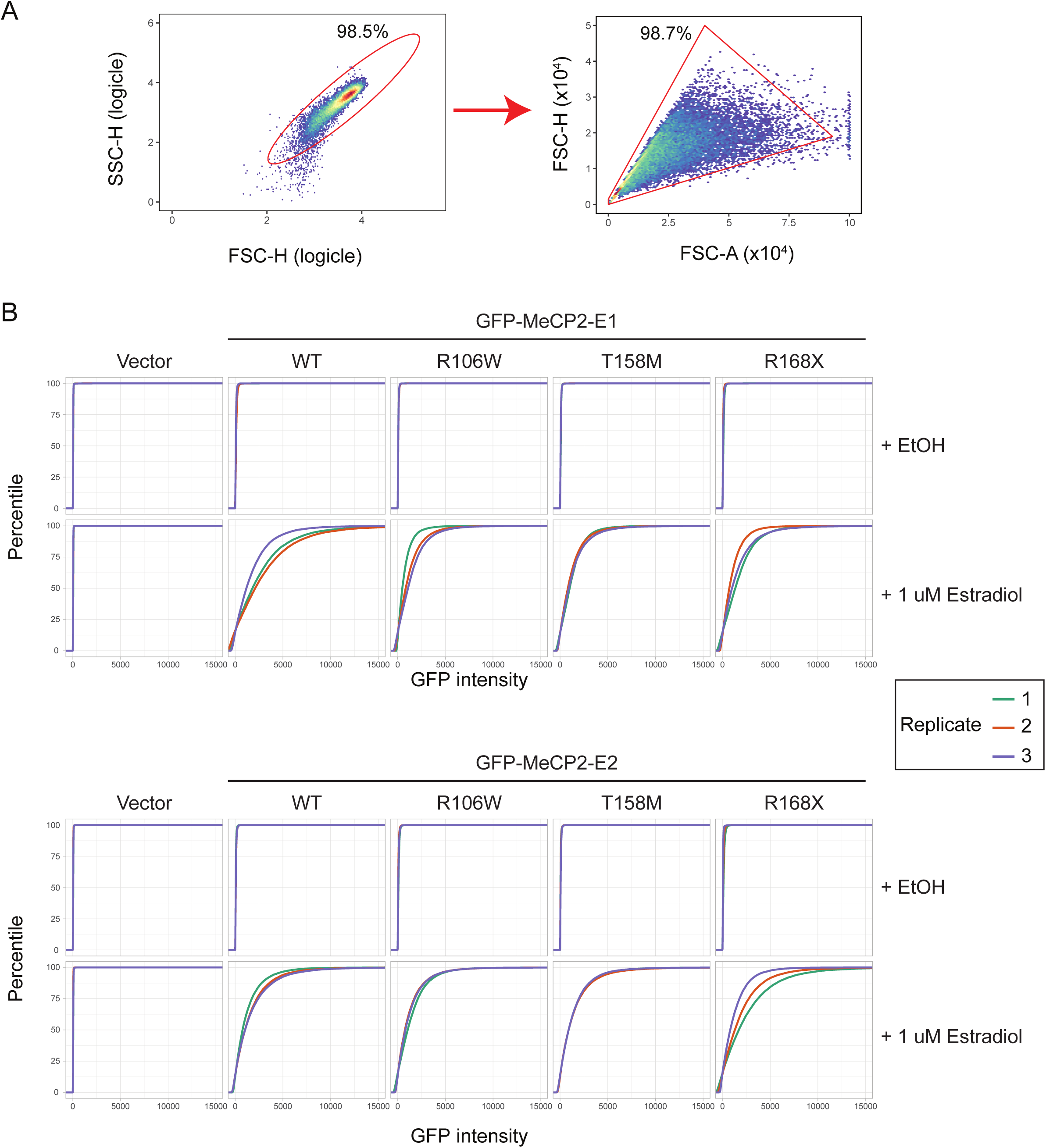
MeCP2 protein level quantification by GFP flow cytometry. A) Gating strategy used for GFP analysis by flow cytometry on live cells. Events were gated by FSC-H vs SSC-H to remove debris, then to remove possible doublet events by FSC-A vs FSC-H. B) Quantified signal from flow cytometry against GFP tag in live cells expressing GFP-MeCP2 constructs after 1 hour of induction with 1 µM estradiol, or ethanol vehicle. This figure shows the flow cytometry performed in Fig. 2c, displayed as empirical cumulative distribution plots.

**Supplementary Figure 6.**
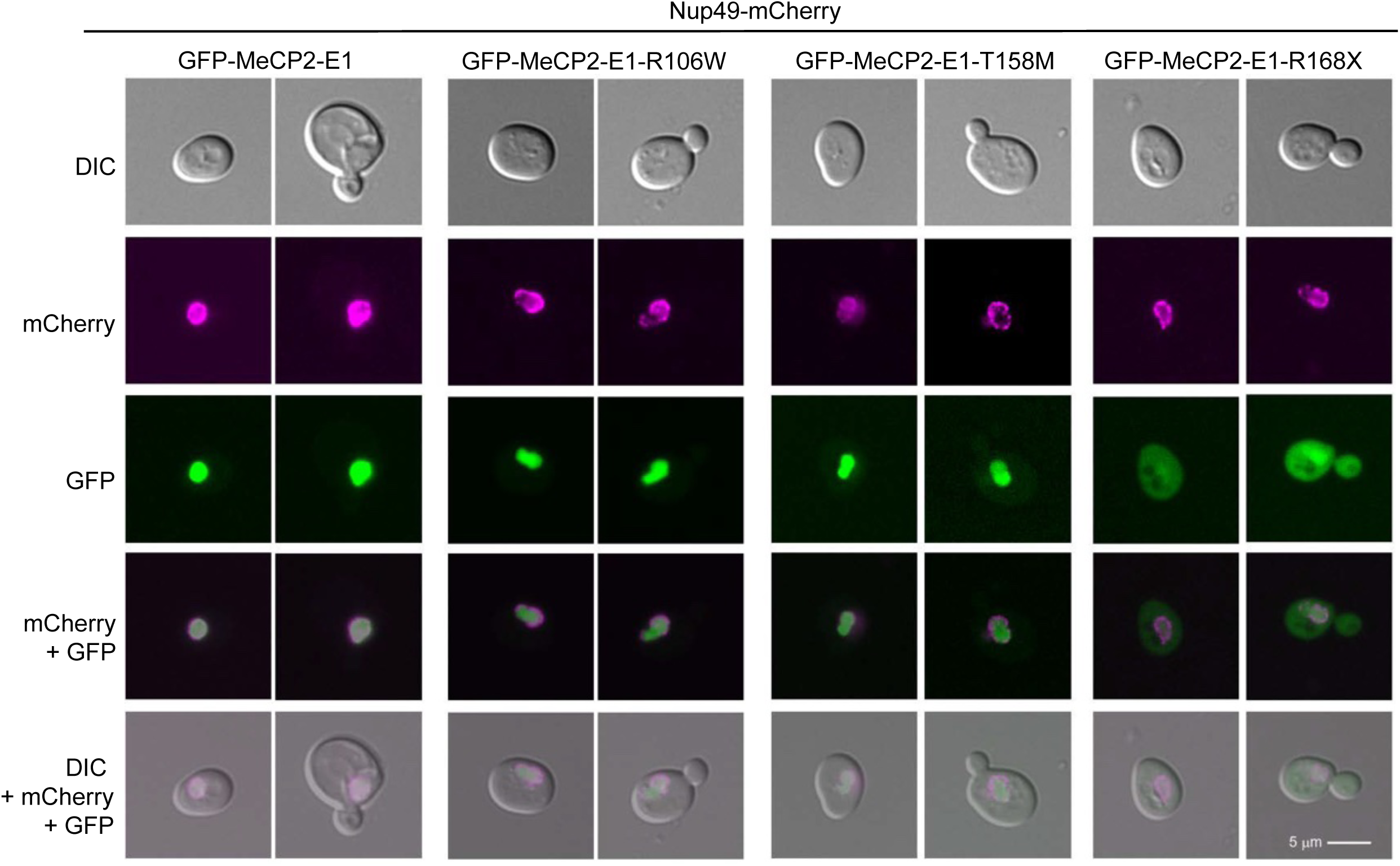
The R168X mutation disrupted the nuclear localization of GFP-MeCP2-E1. Fluorescent microscopy was performed as in Fig. 2d, note that Nup49-mCherry signal is shown as magenta.

**Supplementary Figure 7.**
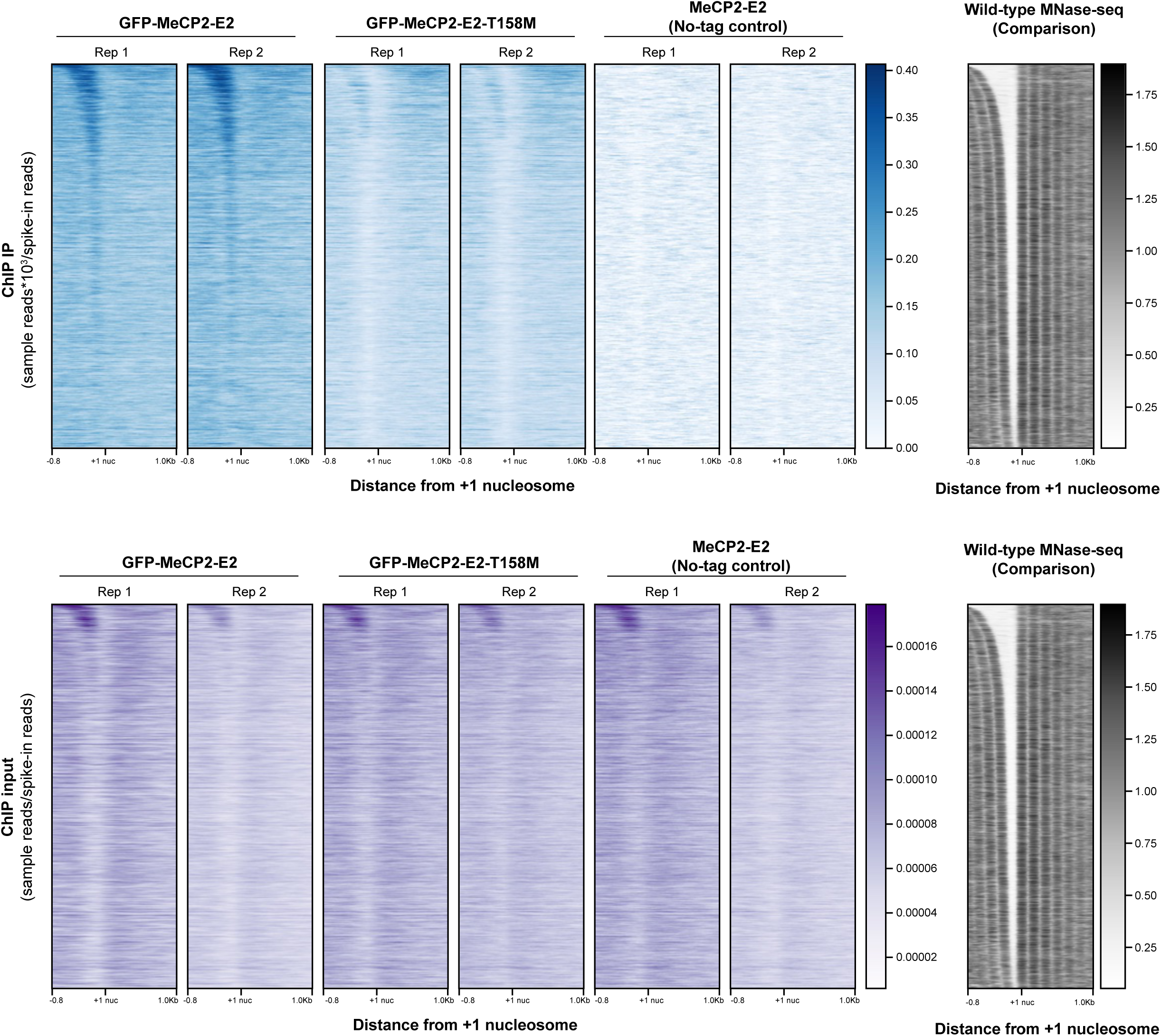
GFP-MeCP2-E2 ChIP-seq data as separate IP and input signals. Two biological replicates are shown. ChIP-seq was performed and analyzed as in Fig. 3.

**Supplementary Figure 8.**
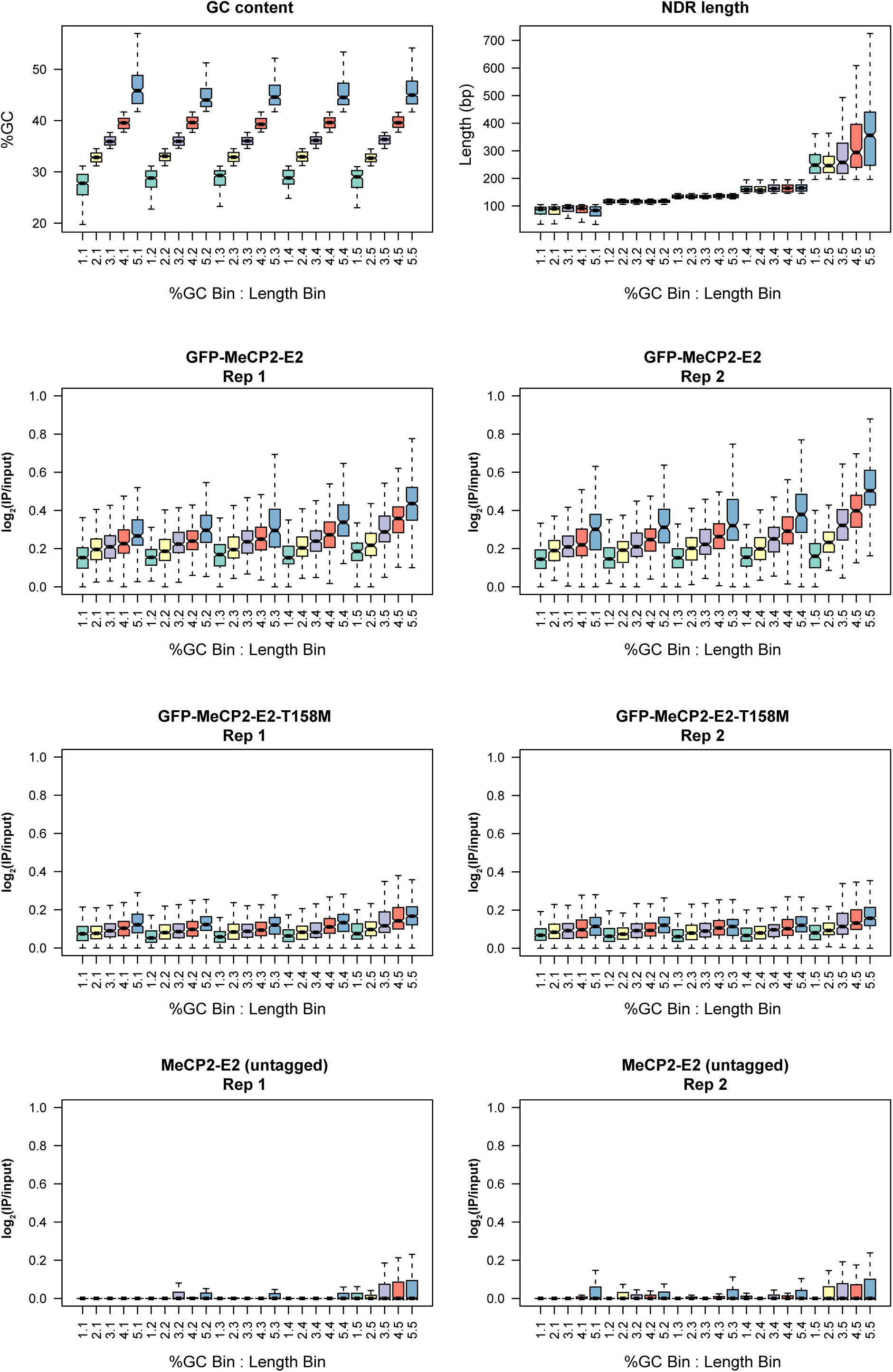
GFP-MeCP2 enrichment within NDRs in ChIP-seq is correlated with %GC. MeCP2 enrichment correlated with %GC, after sorting NDRs into bins by both NDR length^46^ and %GC. Analysis was performed as in Fig. 3b, except that quintiles of GC content were split from quintiles of NDR length. GFP ChIP-seq enrichment (log2 IP/input) is shown for each bin of NDR length and %GC. Boxplots are centered on median signal as described in Methods.

